# Using Constraint-Based Metabolic Modeling to Elucidate Drug-Induced Metabolic Changes in a Cancer Cell Line

**DOI:** 10.1101/2025.05.08.652850

**Authors:** Xavier Benedicto, Åsmund Flobak, Miguel Ponce-de-León, Alfonso Valencia

**Affiliations:** Barcelona Supercomputing Center (BSC); Department of Clinical and Molecular Medicine, Norwegian University of Science and Technology (NTNU), Trondheim, Norway; The Cancer Clinic, St Olav’s University Hospital, Trondheim, Norway; Department of Biotechnology and Nanomedicine, SINTEF Industry, Trondheim, Norway; Catalan Institute for Research and Advanced Studies (ICREA), Barcelona, Spain

**Keywords:** cancer metabolism, drug synergy, constraint-based modeling, systems biology

## Abstract

Cancer cells frequently reprogram their metabolism to support growth and survival, making metabolic pathways attractive targets for therapy. In this study, we investigated the metabolic effects of three kinase inhibitors and their synergistic combinations in the gastric cancer cell line AGS using genome-scale metabolic models and transcriptomic profiling. We applied the TIDE (Tasks Inferred from Differential Expression) algorithm to infer pathway activity changes and introduced TIDE-essential, a new variant that focuses on task-essential genes, enhancing results robustness. Our results revealed widespread down-regulation of biosynthetic pathways, particularly in amino acid and nucleotide metabolism. Combinatorial treatments induced condition-specific metabolic alterations, including strong synergistic effects in the PI3Ki-MEKi condition affecting ornithine and polyamine biosynthesis. These metabolic shifts provide insight into drug synergy mechanisms and highlight potential therapeutic vulnerabilities. To support reproducibility, we developed an open-source Python package, MTEApy, implementing both TIDE frameworks.

## 1. Introduction

Cancer comprises a heterogeneous group of diseases driven by diverse genetic and epigenetic alterations that promote malignant phenotypes such as uncontrolled proliferation, resistance to cell death, and immune evasion [1,2]. These alterations frequently disrupt key signalling pathways that regulate proliferation and apoptosis, leading to sustained growth signals and impaired cell death mechanisms [2]. Signalling deregulation also drives downstream metabolic changes. Cancer cells reprogram their metabolism to meet the demands of rapid growth by producing energy and biosynthetic precursors [3,4]. Pathways such as PI3K/AKT/mTOR, MAPK, and TAK1/NF-*κ*B contribute to this metabolic rewiring and play central roles in oncogenesis [5,6]. High-throughput technologies have transformed our understanding of cancer by providing detailed molecular profiles [7]. However, the complexity of cancer progression and treatment response remains only partially understood. Bioinformatic methods—such as differential expression, over-representation, and gene set enrichment analyses—have been widely used to identify changes in gene activity after perturbations [8,9]. While informative, these approaches are typically descriptive and do not simulate underlying cellular dynamics [10].

Mathematical models can help bridge the gap between molecular interactions and phenotypic outcomes by providing mechanistic insights [11]. Boolean models of cell signalling have been developed for various cancer types and used to predict cell fate and the effects of perturbations such as drug treatments [12,13]. These models have also been tailored with omic data to improve predictions of treatment outcomes [14] and to identify synergistic drug combinations across different cancers [15,16]. In the context of cancer metabolism, genome-scale metabolic models (GEMs) have been reconstructed for humans [17–19] and model organisms such as rats [20]. Constraint-based methods [21] have enabled the integration of omic data into these models to generate context-specific GEMs (CS-GEM) [22–24]. CS-GEM have been applied to cancer cells [25] to investigate metabolic reprogramming [26], identify potential therapeutic targets [27,28], and screen for anti-metabolites [29]. However, the construction of CS-GEM remains challenging due to methodological inconsistencies and a lack of consensus on best practices [30,31]. To address these limitations, alternative approaches such as genetic minimal cut sets [32] have been employed to predict synthetic lethal interactions [33,34]. Additionally, constraint-based algorithms like TIDE (Tasks Inferred from Differential Expression) [35] and CellFie [36] have been developed to infer pathway activity directly from gene expression data, without the need to construct a full CSMM.

In previous work, Flobak et al. constructed a Boolean model of signalling in the gastric adenocarcinoma cell line AGS and identified synergistic effects of kinase inhibitor combinations, which were validated in vitro [15]. Tsirvouli et al. later expanded on these findings using transcriptomic and phosphoproteomic profiling across time points to dissect the mechanisms of synergy [37]. This work aimed to characterise the metabolic alterations induced by three kinase inhibitors and their synergistic combinations in AGS cells. We profiled gene expression changes following drug treatments and used constraint-based modelling to investigate their metabolic effects. Specifically, we applied the TIDE algorithm [35] to infer changes in metabolic pathway activity and proposed a variant named TIDE-essential, which focuses on essential genes without relying on flux assumptions. To quantify synergy at the metabolic level, we introduced a scoring scheme that compares the effects of combination treatments with those of individual drugs. This enabled us to identify metabolic processes specifically altered by drug synergies. We implemented both TIDE and TIDE-essential in an open-source Python package and command-line tool, MTEApy, to facilitate broader use. Our approach provides a framework for investigating drug-induced metabolic rewiring and offers insights into the mechanisms of synergy in targeted cancer therapies.

## 2. Results

### 2.1. AGS cells treated with kinase inhibitors show larger numbers of up-regulated than down-regulated genes

Kinase inhibitors are known to down-regulate pathways involved in proliferation and survival while up-regulating stress response and compensatory mechanisms. Furthermore, combining kinase inhibitors can produce synergistic effects that are absent with individual drugs. In this study, we investigated the functional impact of synergistic drug combinations on various cellular processes, with a particular emphasis on cellular metabolism. We performed a transcriptomic analysis of AGS cells treated with the kinase TAK1, MEK, and PI3K inhibitors (TAKi, MEKi, PI3Ki) and two PI3Ki combinations (PI3Ki-TAKi and PI3Ki-MEKi) and without inhibitors (control condition). We sequenced the transcriptome of AGS cells for the different conditions and identified differentially expressed genes (DEGs) using a standard pipeline (see Materials and Methods). Results showed that the number of DEGs varied across treatment conditions, with an average of ∼2000 DEGs per condition (see Fig. 1A-B left panels and Supplementary Table S1). We observed a larger number of up-regulated or over-expressed genes (∼1200) than down-regulated (∼700) across all treatment conditions. We focused specifically on metabolic genes and found that the patterns remain, suggesting that drugs induce the activity of different metabolic processes (see Fig. 1A-B right panels). The analysis also reveals that MEKi induces the most significant transcriptional changes among individual treatments, followed by TAKi and PI3Ki (see Fig. 1A). Regarding the combinatorial treatments, in PI3Ki-TAKi the number of DEGs is similar to the one observed for TAKi. Nonetheless, the number of DEGs in PI3Ki-MEKi was mildly higher than those reported in either PI3Ki or MEKi, suggesting a possible synergetic effect (see Fig. 1A).

**Figure 1.**
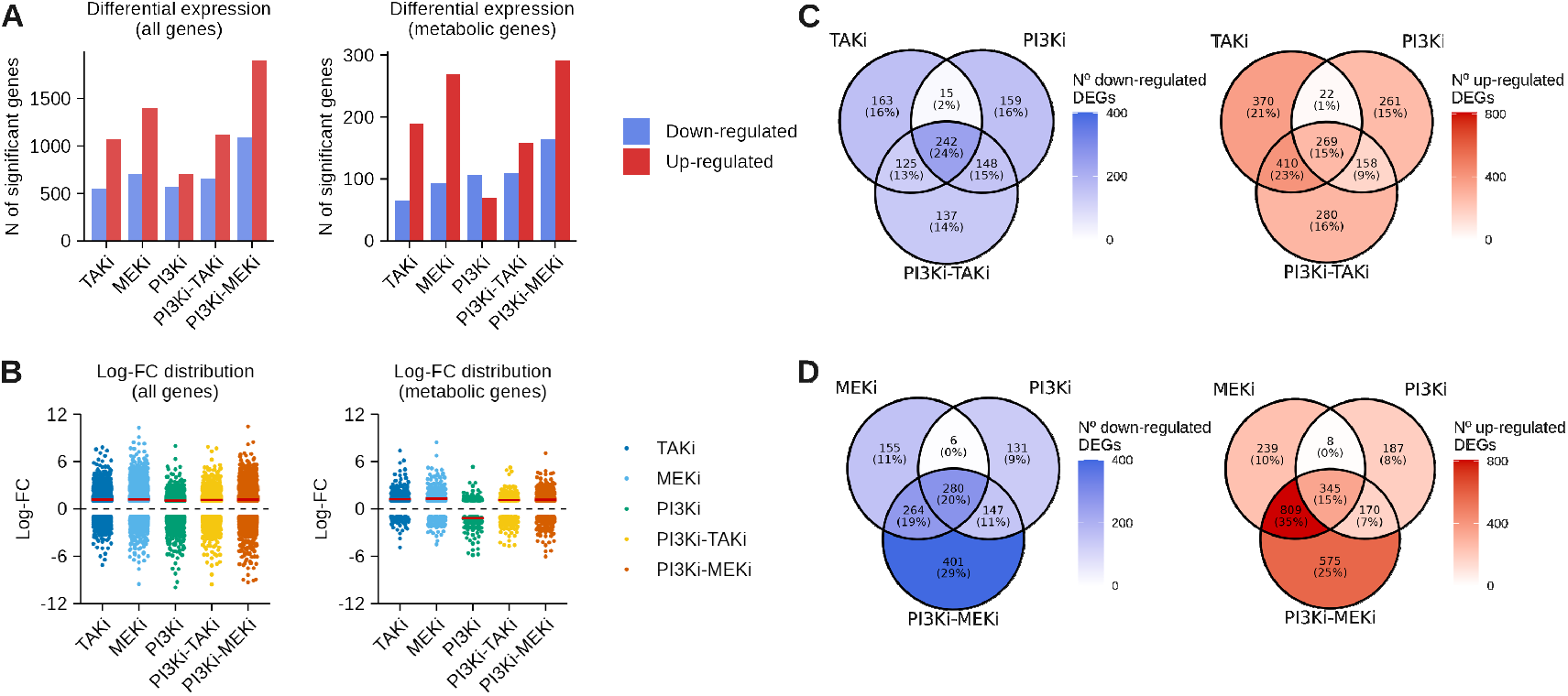
Differential expression analysis on AGS cells after drug perturbations reveals significant shifts. A) Bar charts indicating the number of differentially expressed genes (DEGs) up- and down-regulated per treatment condition for all genes (left panel) and genes within the metabolic model Human-GEM (right panel). Significance was set using an adjusted p-value of less than 0.05, and an absolute log-FC greater than 1. B) Violin plots showing the distribution of DEGs across treatments for all genes (left panel) and genes within the metabolic model Human-GEM (right panel). The median is indicated using a red, horizontal line. C) Venn diagrams for up- (left panel) and down-regulated (right panel) DEGs for the TAKi, PI3Ki and PI3Ki-TAKi treatments. D) Venn diagrams for up- (left panel) and down-regulated (right panel) DEGs for the MEKi, PI3Ki and PI3Ki-MEKi treatments. Darker colours indicate a higher number of DEGs.

We assessed the similarity of up- and down-regulated DEGs across treatment conditions using the Jaccard Index (JI). Shared DEGs may reflect common transcriptional responses, while combination-specific DEGs may suggest drug-specific mechanisms; in particular, changes only observed in the combinatorial treatments may indicate potential synergistic effects. The analysis shows that PI3Ki-TAKi predominantly exhibits an additive effect, with only a small proportion (∼15%) of unique DEGs not observed in single treatments (see Fig. 1C and Supplementary Fig. 6). In contrast, PI3Ki-MEKi demonstrates potentially stronger synergistic effects, evidenced by a larger number of DEGs and a higher proportion (∼25%) of unique DEGs (see Fig. 1D and Supplementary Fig. S6). These unique DEGs may represent distinct pathways activated by the combined effect of drugs and thus may provide insight into the mechanisms of synergies.

To explore the functional implications of the observed gene expression changes, we performed a Gene Set Enrichment Analysis (see Materials and Methods for details). The analysis revealed no strong bias in gene set regulation, with ∼55% down-regulated and 45% up-regulated gene sets on average across conditions (Fig. 2A-B). An exception was observed for PI3Ki-TAKi, which exhibited a significantly larger proportion of down-regulated gene sets (Fig. 2A). Interestingly, this pattern contrasts with the overall gene expression changes, where up-regulated genes predominated across conditions (Fig. 2A). Among individual treatments, MEKi induced the most significant perturbation in biological processes (*n* = 142), followed by TAKi (*n* = 74) and PI3Ki (*n* = 40), consistent with the trends observed in the differentially expressed genes (DEGs).

**Figure 2.**
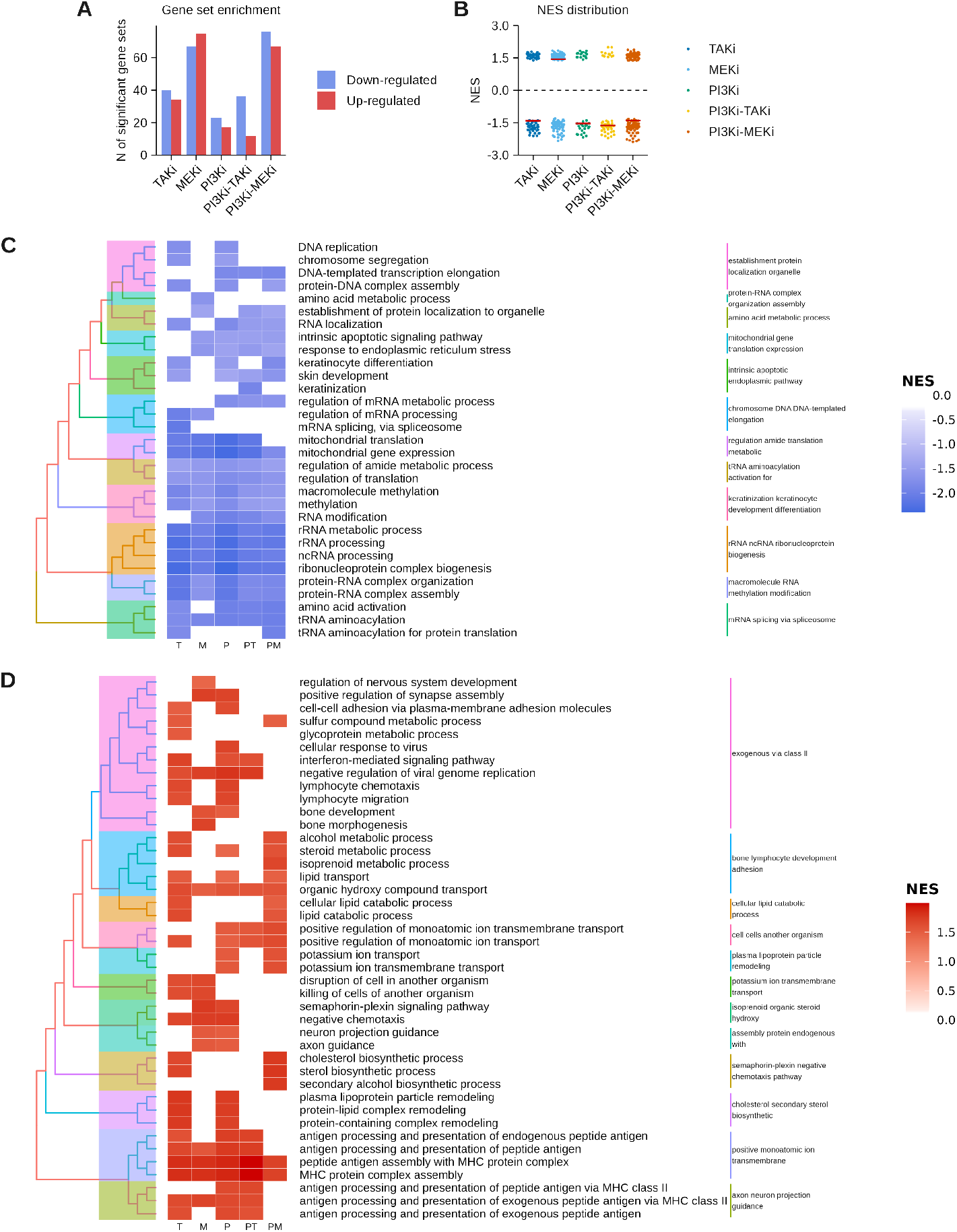
Gene set enrichment analysis summary. A) Bar chart indicating the number of significant gene sets up- and down-regulated per treatment condition. Significance was set using an adjusted p-value of less than 0.05, and an absolute Normalised Enrichment Score (NES) greater than 1. B) Violin plot showing the distribution of DEGs across treatments for all genes. The median is indicated using a red, horizontal line. C) and D) Heatmaps of the top 15 significant gene sets in terms of significance and NES across all treatment conditions for up- and down-regulated gene sets, respectively. Bluer colours in the heatmap indicate lower, negative NES values, while redder colours indicate higher, positive NES values. White cells indicate that the specific gene set in a given condition was not statistically significant. T, M and P correspond to treatments TAKi, MEKi, and PI3Ki, respectively; PT and PM refer to combinations.

We first focus on those processes that are found to be significantly affected in all the conditions. The results show down-regulation of rRNA and ncRNA ribonucleotide biogenesis, rRNA-protein complex organisation and tRNA aminoacylation, suggesting an overall suppression of protein synthesis and translational machinery in the treated cells; down-regulation of mitochondrial gene expression is also found in all the conditions (see Supplementary Fig. 7). Furthermore, we also observed the up-regulation of processes related to the major histocompatibility complex and the transport of organic hydroxy compounds. Cells treated with the individual drugs exhibit similar down-regulation patterns, primarily affecting pathways related to DNA processes, keratinocyte development, and apoptosis (see Fig. 2C). Regarding the up-regulated process, we observed common pathways including metabolism, immune-related processes (e.g., interferon signalling), and condition-specific changes such as lipid metabolism (TAKi) and nervous system development (PI3Ki). Notably, TAKi and PI3Ki display more similar patterns to each other than to MEKi (see Fig. 2D and Supplementary Fig. 7).

We selected the 15 most significant altered gene sets observed in the combinatorial treatments and found that almost all are also altered in the individual treatment counterparts. Nevertheless, we observed some potential synergetic effects in both combinations that include the down-regulation of keratinisation and the regulation of mRNA metabolic process. Next, we focus on those pathways that are significantly altered in the combinatorial treatments but not in the individual drug treatments (see Supplementary Fig. 8). We identified 55 condition-specific gene set alterations in the PI3Ki-MEKi condition, accounting for approximately 40% of all gene sets altered in this condition. For the PI3Ki-TAKi condition, 12 condition-specific gene set alterations were observed, representing around 30% of the significant deregulated gene sets (see Supplementary Fig. 8). We analysed the different gene sets altered in the combinatorial treatments and found that the altered processes represent broad functional categories and do not include specific metabolic processes. Therefore, we propose using a model-driven inference approach to gain deeper insight into the specific metabolic processes altered by the synergistic effects of the drug combinations.

### 2.2. Kinase inhibitors induce down-regulations in key biosynthetic metabolic pathways

Cell growth and proliferation require energy and building blocks that are supplied by different metabolic pathways. Therefore, drugs that affect cell proliferation are likely to have a downstream inhibitory effect on the activity of various metabolic pathways. Herein, we used a constraint-based metabolic modelling approach to better understand the changes in metabolic pathway activity following treatment with kinase inhibitors. Specifically, we used the *Task Inferred from Differential Expression* (TIDE) framework, proposed by Dougherty et al. (2021) [35] to find metabolic pathways significantly altered in the different conditions (see Materials and Methods). We retrieved the most recent version of the genome-scale metabolic reconstruction of human cells, the Human1 model [19]. We also retrieved a list of well-defined metabolic tasks, where each task is defined as a set of input or source metabolites and a set of output or sink metabolites[36]. We manually curated several metabolic tasks to make them compatible with the Human1 model (see Materials and Methods and Supplementary Table S2 for further details).

We ran TIDE on all the conditions using the Human-GEM model and a list of 189 metabolic tasks. The results showed that, on average, ∼50 metabolic tasks were significantly altered in each treatment condition. A total of 30, 17, and 77 significant metabolic tasks were identified in the individual treatments TAKi, MEKi, and PI3Ki; whereas 55 and 66 were identified in the combinatorial treatments PI3Ki-TAKi and PI3Ki-MEKi, respectively (see Fig. 3A and Supplementary Table S3). We observed a trend towards down-regulation of metabolic tasks, with ∼5 and ∼40 up- and down-regulated metabolic tasks, respectively, on average across all treatment conditions (see Fig. 3A). This pattern contrasted with the differential expression results, where we observed more up-regulated DEGs (see Fig. 1A) and is similar to the trends exhibited by the GSEA results (see Fig. 2A).

**Figure 3.**
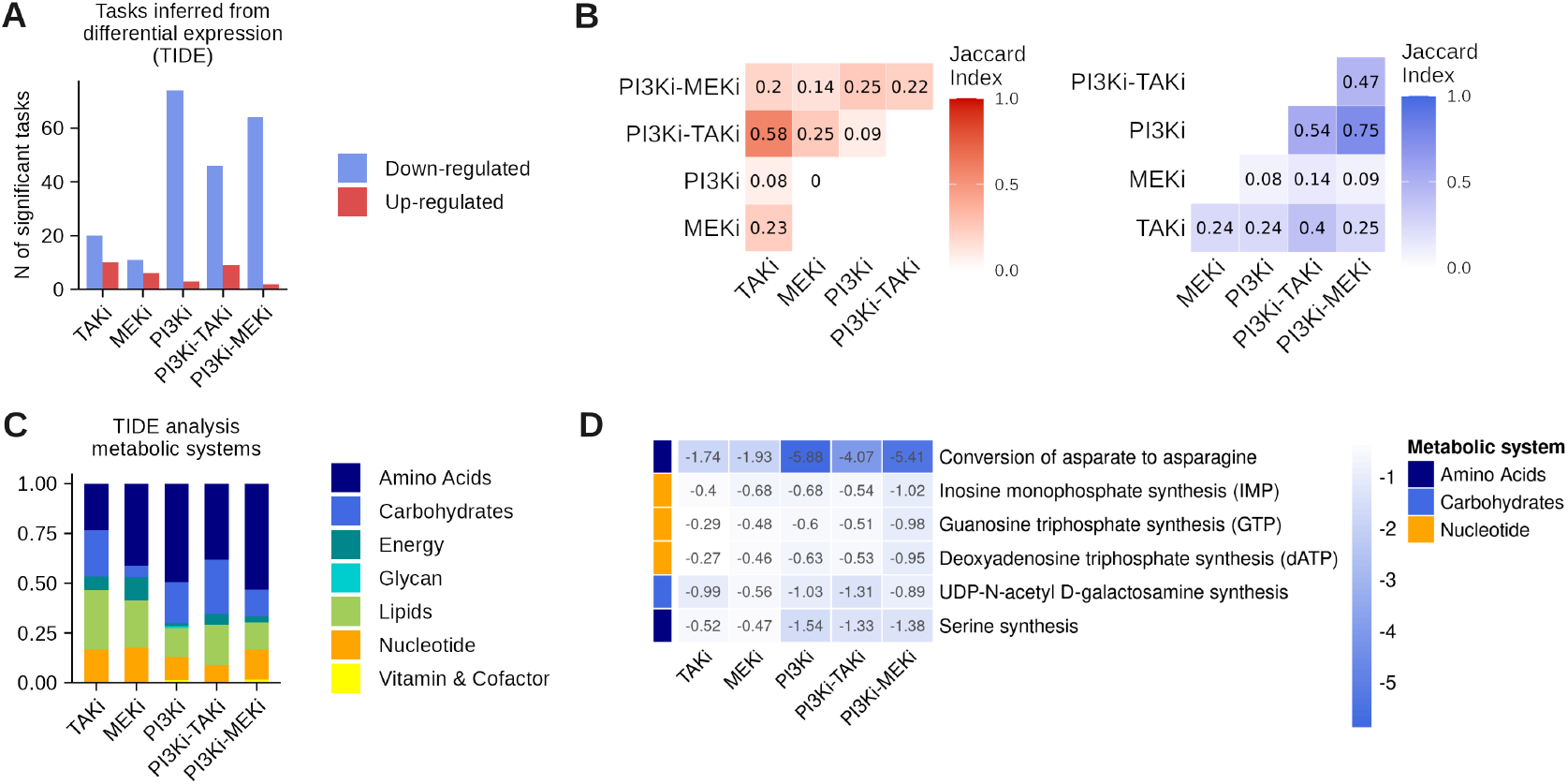
Significant metabolic shifts were identified in AGS cells after drug treatment. A) Bar chart indicating the number of significant metabolic scores up- and down-regulated per treatment condition. Significance was set using a p-value of less than 0.025. B) Jaccard index correlation plots between up-regulated (left panel) and down-regulated (right panel) metabolic tasks across treatment conditions. Darker hues indicate higher correlation values. C) Bar chart indicating the proportions of metabolic systems altered in each condition. D) Heatmap of shared metabolic pathways significantly altered across all the treatments. Darker hues indicate lower, negative metabolic scores. The metabolic system is indicated on the colour strip to the left of the rows.

We assessed the similarities between treatment conditions using the Jaccard Index for the up and down-regulated metabolic perturbations (see Fig. 3B). The highest JI_*up*_ was reported between the two conditions that identified the most up-regulated perturbations, TAKi and PI3Ki-TAKi, while other conditions had less up-regulated perturbations and therefore, lower JI_*up*_ values. This result indicates that TAK1 inhibition triggers the upregulation of some metabolic pathways that are also up-regulated in the combinatorial treatment with PI3Ki. Regarding down-regulated perturbations, PI3Ki and PI3Ki-MEKi displayed the highest similarity values, followed by PI3Ki and PI3Ki-TAKi.

In general, metabolic changes were mainly observed in pathways related to amino acids, carbohydrates, lipids, and nucleotide metabolism (see Fig. 3C). We first analysed pathways affected in all the conditions and found that several metabolic tasks, such as the synthesis and conversion of some amino acids and the production of some nucleotides and deoxynucleotides, were consistently downregulated across all conditions (see Fig. 3D). The results also showed that the conversion of aspartate to asparagine and serine synthesis were down-regulated across all treatment conditions. Notably, we observed that in PI3Ki, or any of its combinations, more amino acid perturbations were identified than in any other single treatment.

### 2.3. Synergistic metabolic shifts in AGS cells were identified following combinatorial drug treatments

The TIDE results showed that most of the metabolic changes identified in the individual treatments are also observed in the combinatorial treatments. However, we also found several examples of metabolic processes exhibiting changes only in the combinatorial treatments. We focused on these unique metabolic perturbations, since they may represent synergetic effects. We introduced a scoring system to quantify weak and strong synergistic effects in the regulation of metabolic pathways. We considered weak synergies cases where the metabolic score in combinatorial treatments is higher/lower than the maximum/minimum observed in their single-drug conditions. Conversely, we defined strong synergies as cases where individual drugs show no effect on the expression of metabolic processes, while the combinatorial treatment significantly alters the process. Noting *A* and *B* as two individual treatments, *AB* as their combination, *t* as a given metabolic task, and *MS*(*A, t*) as the corresponding metabolic score of task *t* in the condition *A*, we defined weak synergies as follows:

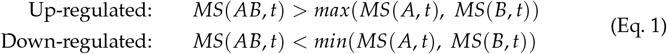

and a strong synergy is defined by the following expression:

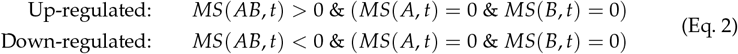

In the PI3Ki-TAKi condition, we found that 14 out of 55 significantly altered metabolic processes exhibit some kind of synergistic effect (see Fig. 4A and Supplementary Table 4). On the other hand, the PI3Ki-MEKi combination exhibited a higher value of synergetic perturbation, including 49 out of 66 significantly affected pathways (see Fig. 4A). Interestingly, both non-synergistic and synergistic perturbations observed in the PI3Ki-TAKi and PI3Ki-MEKi treatments show a bias towards down-regulation or inhibition of metabolic processes (see Supplementary Tables 3 and 4). In addition, there were no differences reported between non-synergistic and synergistic metabolic scores in PI3Ki-TAKi (Mann-Whitney U test p-value = 0.363), but some slightly significant differences were reported between scores in PI3Ki-MEKi (p-value = 0.043) (see Fig. 4B). Most of the synergistic perturbations in PI3Ki-TAKi included pathways related to the lipid (*n* = 5) and amino acid (*n* = 4) metabolism (see Fig. 4C-D). In PI3Ki-MEKi, however, apart from the aforementioned systems (*n* = 8 and *n* = 25, respectively), synergistic perturbations were also observed in the nucleotide (*n* = 10) and carbohydrate (*n* = 5) metabolism, among others (see Fig. 4C-D).

**Figure 4.**
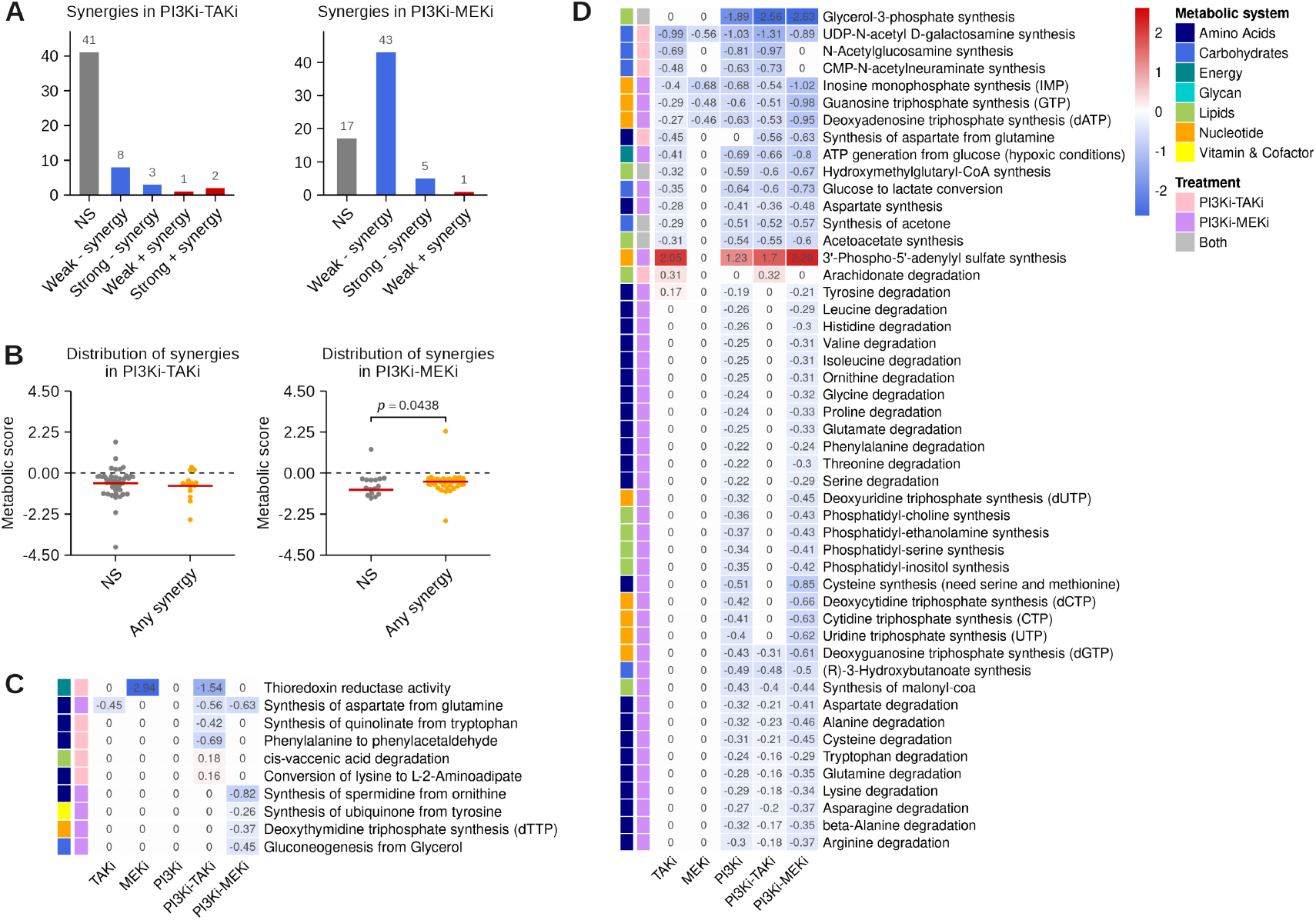
Synergistic effects over AGS metabolism trend towards down-regulation. A) Bar chart indicating the number of significant, synergistic metabolic scores for PI3Ki-TAKi and PI3Ki-MEKi. Synergy type is indicated in blue for down-regulated scores and in red for up-regulated scores. Non-synergistic scores (NS) are indicated in grey. B) Distribution plots comparing the distributions of NS and synergistic scores regardless of their type. The mean is indicated with a horizontal red line. C) Heatmap of strong synergies for either PI3Ki-TAKi or PI3Ki-MEKi. Scores for all treatment conditions are shown for comparison. Coloured bars indicate the metabolic system (leftmost) and the combinatorial treatment to which the synergy belongs. Panel D) shows a heatmap of weak synergies for either PI3Ki-TAKi or PI3Ki-MEKi. Scores for all treatment conditions are shown for comparison. Coloured bars indicate the metabolic system (leftmost) and the combinatorial treatment to which the synergy belongs.

Strong synergies regarding the PI3Ki-TAKi treatment involved the down-regulation of the presence of the thioredoxin system through the thioredoxin reductase activity and some alterations in the phenylalanine and tryptophan metabolism. Pathways related to lysine metabolism and degradation of cis-vaccenic acid were slightly up-regulated in PI3Ki-TAKi (see Fig. 4C). Interestingly, all strong synergies involved in PI3Ki-MEKi were found to be down-regulated, and included a wide range of processes from ornithine and aspartate biosynthesis, gluconeogenesis, and deoxynucleotides to ubiquinone 10 (Q10) biosynthesis. Regarding weak synergies, more perturbations were observed in PI3Ki-MEKi than in PI3Ki-TAKi, and only 2 were identified as up-regulated across treatment conditions. Interestingly, four weak synergies were common to PI3Ki-TAKi and PI3Ki-MEKi conditions: glycerol-3-phosphate synthesis, hydroxymethylglutaryl-CoA synthesis, and acetone and acetoacetate synthesis.

### 2.4. The TIDE-essential framework identifies new metabolic perturbations and synergistic effects in AGS metabolism

In this section, we introduced a variation of the TIDE algorithm, named TIDE-essential, that aims to extend the framework (see Materials and Methods for more details). In the TIDE algorithm, each metabolic task, defined by a set of input and output metabolites, is first associated with a feasible flux distribution. Due to the presence of isozymes or alternative pathways, a metabolic task may have multiple flux distributions that fulfil its requirements. TIDE resolves this by using Parsimonious Flux Balance Analysis (pFBA) to compute a single flux distribution that minimises total flux while satisfying the task constraints (see Materials and Methods). The activity change of the pathway is then estimated by projecting gene expression changes (log-fold changes) onto the flux-carrying reactions of this reference solution. However, because this flux distribution is computed before integrating gene expression data, it may not reflect the biologically relevant flux patterns under perturbed conditions.

To address this potential bias, we propose a more conservative variation of the TIDE framework that does not rely on a precomputed flux distribution. Instead, it evaluates task activity based solely on the set of essential genes required to complete each metabolic task (Fig. 5A). These essential genes are identified through *in-silico* single-gene knockout analysis and represent the minimal set necessary to produce the output metabolites from the specified inputs (see Materials and Methods). The expression changes of the essential genes are then used to assess task activity, with statistical significance evaluated using the same procedure as in TIDE, by computing an empirical null distribution of expression changes to determine whether the observed shift is significant. This essential-gene-based formulation of TIDE enhances robustness in the analysis of metabolic perturbations for two main reasons: (i) it eliminates the dependency on pFBA-derived flux distributions to identify task-associated reactions, and (ii) it operates directly at the gene level, increasing interpretability and relevance in transcriptomics-based analyses.

**Figure 5.**
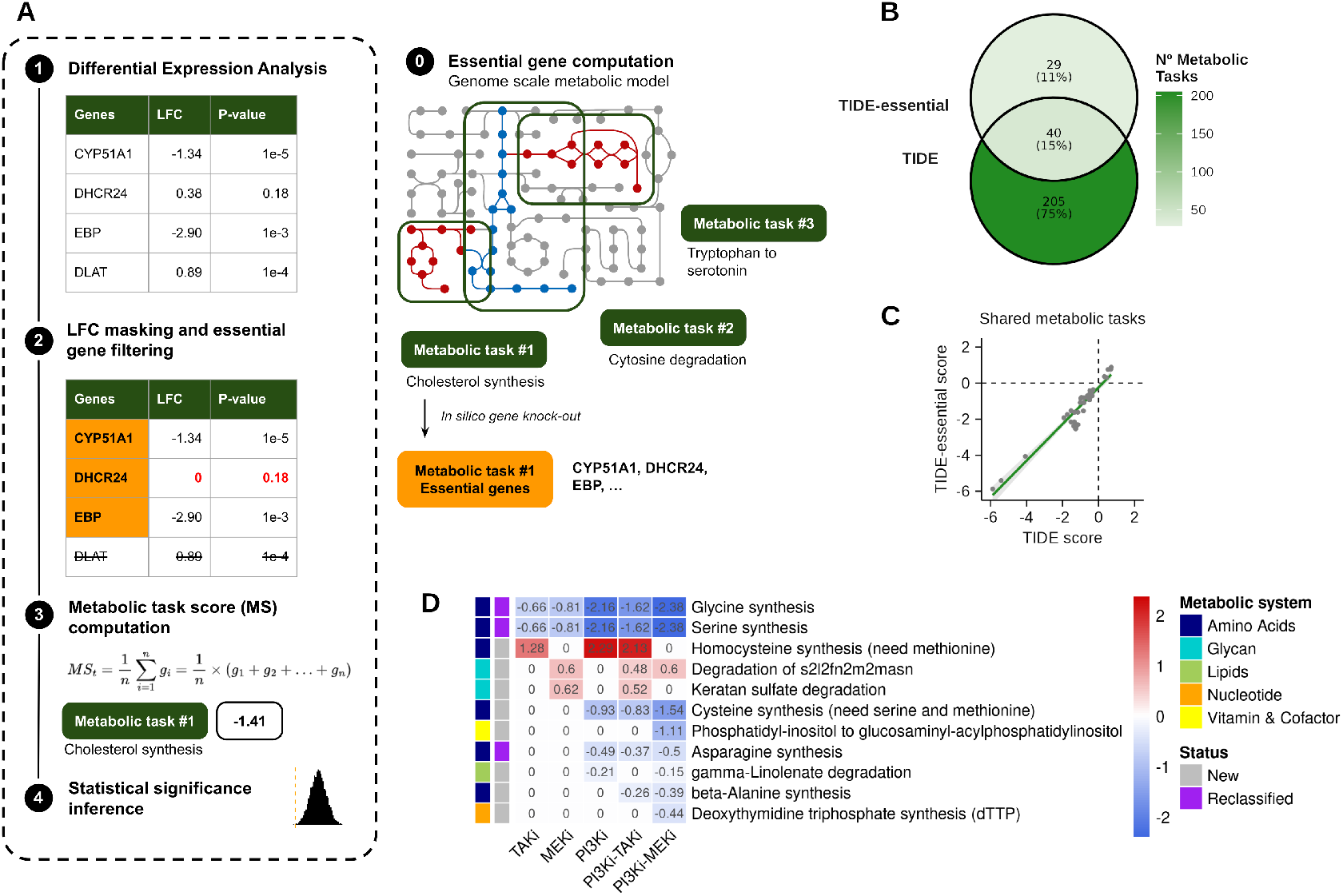
New metabolic perturbations and synergistic effects in AGS metabolism can be inferred using TIDE-essential. A) Schematic overview of the TIDE-essential framework. B) Venn diagram showing the overlap of significant metabolic tasks between the TIDE and TIDE-essential framework across treatment conditions. C) Scatter plot of the scores of the 40 common metabolic tasks between the TIDE and TIDE-essential frameworks. The orange line indicates the fitted linear model, with a Spearman ranked correlation value of 0.93 (p-value *<* 1−10). Treatment conditions are indicated by different shapes. D) Heatmap of the new and reclassified metabolic perturbations identified by TIDE-essential. Redder hues indicate higher, positive metabolic scores and bluer hues indicate lower, negative values. The metabolic system and the origin of the showcased metabolic tasks are indicated on the colour strips to the left of the rows.

We applied TIDE-essential to the same metabolic tasks and treatment conditions as in the original TIDE analysis. TIDE-essential identified a total of 69 metabolic tasks, of which 40 overlapped with those detected by TIDE (Fig. 5B). The metabolic scores derived from both methods were highly correlated (Spearman’s *ρ* = 0.93; Fig. 5C), indicating overall agreement between the two approaches. Although TIDE generally identified a higher number of altered tasks across conditions, for instance, in the PI3Ki treatment, TIDE detected 70 unique tasks compared to 3 by TIDE-E—TIDE-essential uncovered a subset of tasks not detected by TIDE (see Table 1 and Supplementary Table 5). This was particularly evident in the MEKi condition, where TIDE-E identified 9 unique tasks versus 10 by TIDE, with 7 tasks in common. These findings suggest that TIDE-essential can complement TIDE by capturing additional condition-specific alterations.

**Table 1.** Tasks Detected by Condition.

| Altered Tasks | Treatment |  |  |  |  |
| --- | --- | --- | --- | --- | --- |
|  | TAKi | MEKi | PI3Ki | PI3Ki-MEKi | PI3Ki-TAKi |
| Unique TIDE | 24 | 10 | 70 | 52 | 49 |
| Unique TIDE-E | 8 | 9 | 3 | 4 | 5 |
| Common | 6 | 7 | 7 | 14 | 6 |
| Total TIDE | 25 | 17 | 77 | 66 | 55 |
| Total TIDE-E | 14 | 15 | 10 | 18 | 11 |
| **Total** | **38** | **26** | **80** | **70** | **60** |

TIDE-essential identified 11 and 18 significant metabolic perturbations in the PI3Ki–TAKi and PI3Ki–MEKi combination treatments, respectively, revealing a consistent trend toward down-regulation, in line with previous observations (Fig. 5D). Notably, in the PI3Ki–TAKi condition, TIDE-E detected five metabolic tasks not reported by TIDE, including three associated with strong synergy. Among these, beta-alanine synthesis was down-regulated, while glycan degradation was broadly up-regulated. In contrast, homocysteine synthesis was up-regulated and cysteine synthesis down-regulated, though neither was linked to synergistic effects (Supplementary Table 5). In the PI3Ki–MEKi condition, TIDE-E uncovered four additional perturbations, two of which displayed strong synergy, namely, the down-regulation of beta-alanine synthesis and a phosphatidylinositol conversion. Furthermore, glycan degradation was up-regulated and gamma-linolenate degradation was downregulated, although these changes were not classified as synergistic. Finally, TIDE-essential reclassified three amino acid synthesis tasks—previously labelled as non-synergistic by TIDE—as weakly synergistic.

## 3. Discussion

In this study, we explored the functional and metabolic effects of individual kinase inhibitors and their synergistic combinations in a cancer cell line. We observed a general trend of gene overexpression across all treatment conditions. Although individual gene expression levels showed an overall trend of overexpression, pathway-level analysis revealed the opposite effect, suggesting that transcriptional changes may be part of a broader regulatory compensation mechanism. This underscores the importance of integrating gene-level data with pathway and metabolic models to reveal biologically meaningful responses.

GSEA results showed that several gene sets related to metabolic processes were down-regulated after treating the cells. However, these results lacked pathway-level resolution, making it difficult to infer precise functional consequences. For instance, the gene set associated with amino acid metabolism identified in GSEA does not specify which particular amino acids are affected, nor does it clarify whether the altered pathway relates to biosynthesis or degradation. Similar results have been found for other functional categories, such as lipid and nucleotide metabolism. In contrast, constraint-based modelling enables the assessment of metabolic tasks at a mechanistic level, revealing pathway-specific alterations.

We integrated differential gene expression into the genome-scale metabolic model of human cells using the TIDE algorithm to identify metabolic pathways altered under different treatment conditions. The results revealed significant down-regulation of several metabolic pathways, particularly those involved in nucleotide synthesis, amino acid metabolism, and lipid biosynthesis. We also developed TIDE-essential, a variant of TIDE that uses only task-essential genes and bypasses the need for flux distributions when estimating pathway activity. Although TIDE identified higher numbers of altered tasks across conditions, we found that TIDE-essential uncovers several altered metabolic tasks, not detected by TIDE, showing that the two algorithms are complementary.

Using this novel methodology, we have identified several essential metabolic pathways relating to modifications that could be produced by drugs and drug combinations at the level of metabolism. Specifically, the nucleotide synthesis (IMP, GTP and dATP) were down-regulated under all treatment conditions, but especially in the PI3Ki-MEki combination. A reduction in the supply of DNA and RNA building blocks is likely related to the reduction in cell growth observed in AGS cells. These findings align with previous studies reporting reduced nucleotide biosynthesis and downregulation of aminoacyl-tRNA synthetase enzymes under similar conditions [37].

In general, we found that treatments involving PI3Ki consistently exhibited a stronger inhibition of metabolic pathways than the other conditions, especially regarding amino acid metabolism. These results are in agreement with previous studies highlighting the central role of the PI3K/AKT/mTORC1 pathway in cancer metabolism [5].

Notably, the conversion of aspartate to asparagine was found to be the top downregulated metabolic task across all treatment conditions, with the most pronounced effect observed in cells treated with PI3Ki. The PI3K/AKT/mTORC1 signalling pathway is known to positively regulate asparagine production [38], which aligns with our observation of enhanced down-regulation in conditions involving PI3Ki and its combinations. Furthermore, the inhibition of asparagine synthetase has been shown to enhance sensitivity to metabolic stress [38], which may contribute to the observed reduction in AGS cell proliferation.

With the introduction of a methodology to quantify the degree of synergy in metabolic pathways, we have obtained relevant results related to the synergistic effects of drug combinations in cellular metabolism. The results indicated that the PI3Ki-MEKi combination had a more pronounced synergistic effect on metabolic pathways compared to the PI3Ki-TAKi combination, as evidenced by a greater number of synergistic alterations, particularly in pathways involving cofactors and vitamins. This aligns with in-vitro assays indicating that the PI3Ki-MEKi combination is more effective in reducing cell proliferation than the PI3Ki-TAKi combination [15].

The strong synergistic effect observed in PI3Ki-MEKi is likely due to the inhibition of pathways not altered by the individual drugs. For instance, we found that the synthesis of spermidine is only observed down-regulated in PI3Ki-MEKi. Polyamines are essential for DNA stability, cell proliferation, and stress resistance [39], and their depletion has been linked to increased DNA damage and apoptosis [40–42]. We also found that ubiquinone (CoQ) synthesis was strongly down-regulated in PI3Ki-MEKi, specifically through reduced expression of coenzyme Q3 methyltransferase (COQ3). COQ3 expression has recently been reported to be associated with worse patient survival in oesophageal adenocarcinoma [43], thus a down-regulation in this metabolic pathway could potentially hinder cancer progression in AGS cells as well.

The PI3Ki-TAKi condition exhibited less synergetic effects in the metabolic pathways than PI3Ki-MEKi. Nevertheless, we found that PI3Ki-TAKi induced a down-regulation in the thioredoxin pathway, suggesting a strong synergistic effect. Thioredoxin protein levels are implicated in tumour growth and apoptosis inhibition [44], and recently, thioredoxin has been proposed as a therapeutic target in lung cancer for its ability to increase sensitivity to CHK1 inhibitors [45]. We also identified a down-regulation in the synthesis of quinolinate from tryptophan, a pathway that has been associated with cell death and cell cycle arrest in colorectal cancer models [46].

By integrating transcriptomic data using TIDE and TIDE-essential, we provide a systematic framework for uncovering metabolic responses to drug treatments. We identified specific pathways down-regulated exclusively by synergistic combinations by analysing metabolic alterations induced by single drugs and synergistic combinations. These findings offer insight into the molecular basis of drug synergies and reveal metabolic vulnerabilities that could be exploited as novel therapeutic targets. To enhance reproducibility and facilitate adoption, we have implemented the TIDE and TIDE-essential algorithms in the open-source Python package MTEApy, offering an accessible tool for the metabolic modelling community. This framework readily applies to other cancer types and drug combinations, supporting broader applications in precision oncology.

## Materials and Methods

### Experimental setup

#### Cell Culture and Drug Treatments

AGS (human gastric adenocarcinoma, ATCC, Rockville, MD) were grown in Ham’s F12 medium (Invitrogen, Carlsbad, CA) supplemented with 5% fetal calf serum (FCS; Euroclone, Devon, UK) and 10 U/ml penicillin-streptomycin (Invitrogen). AGS cells were seeded in 6-well plates and treated after 24 hours with kinase inhibitors (5Z)-7-oxozeaenol (TAKi), PD0325901 (MEKi), and PI103 (PI3Ki), individually or in combination, for another 24 hours. Chemical inhibitors (5Z)-7-oxozeaenol, PD0325901, and PI-103 (all Merck) were dissolved in DMSO at stock concentrations of 20 mM. Each drug treatment was administered to concentrations previously reported to reduce growth by 50% (GI_50_ concentrations) after 48 hours compared to the vehicle control, dimethyl sulfoxide (DMSO) [15]. Cells were also treated using the combination of these drugs (i.e. PI3Ki-TAKi and PI3Ki-MEKi) that were experimentally confirmed to be synergistic in AGS cells[15]. For the combinatorial treatments, the half concentrations of GI_50_ were used for each drug (see Table 2).

**Table 2.** Description of the kinase inhibitor used.

| Compound | Short name | Target | GI50 |
| --- | --- | --- | --- |
| (5Z)-7-oxozeaenol | TAKi | TAK1 | 0.500 μM |
| PD0325901 | MEKi | MAP2K1/2 | 0.035 μM |
| PI103 | PI3Ki | PI3KCA | 0.700 μM |

#### RNA Sequencing

After treatment, cells were lysed in RNA lysis buffer (Qiagen RLT Plus) and stored at -80°C. RNA extraction was performed using the AllPrep DNA/RNA/miRNA Universal Kit (Qiagen). RNA concentration and integrity were assessed using the Qubit RNA HS Assay and Agilent Bioanalyzer, respectively. Libraries were prepared with the Illumina Stranded mRNA Prep Ligation Kit, purified with AMPure XP beads, quantified via qPCR, and validated on a Bioanalyzer. Sequencing (2×64 bp) was performed on an Illumina HiSeq2000 platform. For each treatment condition and the control, four replicates were performed.

### Bioinformatic analysis

#### RNA-Seq Data Processing and Quantification

FASTQ files were generated with bcl2fastq (v2.20), quality-checked with fastqc (v0.12.1), and preprocessed using fastp (v0.20.1). Transcript abundances were estimated using Salmon (v1.10.3) with the reference Human genome GRCh38. Quality control summaries were generated using MultiQC (v1.23) and gene-level counts were computed using the *tximport* R package (v1.32.0).

#### Differential Expression Analysis

*Differential expression analysis (DEA)* was performed using the *DESeq2* framework (version 1.40.0) [47]. A linear model was fitted for each gene and condition with the different treatments as a variable, correcting for possible confounding effects using the different replicates. The DMSO samples were used as the baseline for contrasts. We applied Benjamini-Hochberg correction for multiple hypothesis testing. A log-FC shrinkage was applied using the *apeglm* normalization method [48]. Differentially expressed genes were defined using an absolute log fold-change threshold of 1 and an adjusted p-value of less than 0.05.

#### Gene set enrichment analysis

Gene set enrichment analysis (GSEA) using Gene Ontology (GO) terms was performed using the *ClusterProfiler* library (version 4.10.0) in R, using the log2-FC ranking of all genes in the study. Significant gene sets were defined using a p-value threshold of less than 0.05 and an absolute Normalised Enrichment Score (NES) greater than 1. Redundancy in GSEA results was managed using the *simplify* function from the *ClusterProfiler* library using a similarity cutoff of 0.75. GSEA results were visualised using *enrichplot* (version 1.24.2).

### 3.1. Genome scale-metabolic modelling

#### Human metabolic model

We retrieved the genome-scale metabolic model Human-GEM version 1.18 from the official repository (https://github.com/SysBioChalmers/Human-GEM). The model accounts for 2897 genes, 11364 metabolic reactions, 1721 exchange fluxes and 8499 metabolites and is the most comprehensive description of human metabolism [19]. We used the list of selected metabolic tasks previously published by Richelle *et al*. (2021) which includes 195 metabolic tasks associated with 7 metabolic systems (energy, nucleotide, carbohydrate, amino acid, lipid, vitamin & cofactor and glycan metabolism).

#### Modelling Metabolic tasks

A metabolic task *t* is defined metabolic task is formally defined as a set of input metabolites 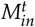 and a set of output metabolites 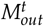 [49]. Each input metabolite *I* is associated with a source reaction 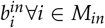, and each output metabolite *j* has an associated sink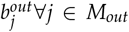. To compute a flux distribution for the task, all exchange reactions are set to zero except those corresponding to the sources and sink reactions associated with the task; then a flux distribution is found by solving and FBA Eq. 6 or pFBA Eq. 7 problem (see Supplementary Material and Methods for further details). Examples include the synthesis of amino acids and nucleotides as well as the generation of energy through glycolysis and the respiratory chain. To implement a metabolic task *t ∈ T* in a model, the exchange reactions are first bounded to zero, and then source and sink boundary reactions are added for the set of inputs and outputs, respectively. Although this list of tasks was defined for the Recon 2.2 model [18], the Human-GEM repository already includes a translated version of these metabolic tasks. A preliminary analysis showed that 9 tasks were inconsistently defined due to incorrect mapping of metabolite identifiers between metabolic Recon2.2 and the Human-GEM. Additionally, we found 18 tasks that resulted in infeasible problems due to problems with bounds for the inputs and the outputs. Inconsistent tasks were manually curated by correcting metabolite identifiers. To curate infeasible tasks, we manually adjusted upper and lower bounds for the input and/or output metabolites or by updating the compartments of some metabolites. After the curation process, 19 metabolic tasks were reconciled to the Human-GEM model, whereas 8 tasks remained either inconsistent (1) or infeasible (7). A summary of the curation processes can be found in Supplementary Table 1. For each consistent metabolic task, pFBA was run to obtain a reference flux distribution.

#### Tasks inferred from differential expression

To investigate changes in metabolic pathway activity, we applied the Tasks Inferred from Differential Expression (TIDE) algorithm [35]. We used the manually curated list of 187 metabolic tasks and their reference flux distributions. Log-fold change (log-FC) values from differential gene expression analysis were mapped onto the flux-carrying reactions associated with each task via their gene–protein–reaction (GPR) rules. In this mapping, each gene was substituted with its corresponding log-FC value. GPR rules were evaluated by interpreting logical AND and OR operators as the *min* and *absmax* functions, respectively, and resolving the resulting expression recursively. Genes with non-significant adjusted p-values were masked by setting their log-FC values to zero before evaluation. The statistical significance of the resulting task scores was assessed through 10,000 permutations. A task was considered significantly altered if its associated p-value was below 0.025, reflecting a two-sided testing strategy.

#### TIDE essential genes

To extend and complement the original TIDE framework, we developed an approach focused on the identification of essential genes required for the execution of curated metabolic tasks. For each task *t*, we performed an *in silico* gene knockout screening across all genes in the metabolic model. Specifically, for each gene *g*, the procedure consisted of the following steps: (1) identifying the set of reactions deactivated by the knockout of *g*; (2) constraining the flux of these reactions to zero; and (3) computing the flux distribution for task *t* using parsimonious Flux Balance Analysis [50] (see Supplementary Material and Methods for further details). If the optimization problem became infeasible, gene *g* was classified as essential for task *t*. This procedure was repeated for all genes and tasks, resulting in a gene-task matrix representing the essential genes for each metabolic task. The metabolic score of a task *t* was computed as the average log-fold change (log-FC) of the essential genes associated with that task. As in the original TIDE framework, log-FC values with non-significant adjusted *p*-values were masked to zero before evaluation, and statistical significance was assessed using 10,000 permutations. A two-sided testing approach was applied, with a significance threshold of *p <* 0.025.

### 3.2. Computational tools

All statistical analyses were conducted using R (version 4.4.1). Figures were generated using the following R libraries: *tidyplots* (version 0.1.2), *ggcorrplot* (version 0.1.4.1), *ggVennDiagram* (version 1.5.2), and *pheatmap* (version 1.0.12). TIDE and TIDE-essential algorithms were implemented in an open-source Python package and command-line tool, MTEApy, built on top of the *COBRAPY* library. The source code is available at: https://github.com/bsc-life/mteapy.

## Acknowledgments

This work has received funding from the Horizon 2020 projects INFORE (ID: 825070) and PerMedCoE (ID: 951773).

## Author Contributions

Conceptualization, X.B., M.P., Å.F., and A.V.; methodology, X.B., Å.F. and M.P.; experimental analysis, Å.F.; software, X.B. and M.P.; in-silico experiments, X.B.; results analysis, X.B and M.P..; writing - original draft preparation, X.B. and M.P.; writing - review and editing, X.B., M.P., Å.F., and A.V.; visualization, X.B.; supervision, M.P. and A.V. All authors have read and agreed to the published version of the manuscript.

## Conflicts of Interest

The authors declare no competing interests.

## Data Availability Statement

AGS cells’ raw RNA sequencing data have been deposited to the GEO repository with the dataset identifier GSE285616. The code used for all presented analyses and all the results is available at https://github.com/bsc-life/ags-paper. The Python library for Metabolic Task Enrichment Analysis (*MTEApy*) is available at https://github.com/bsc-life/mteapy.

## Notes

### Competing Interest Statement

The authors have declared no competing interest.

