## Supplementary Material for "Using Constraint-Based Metabolic Modeling to Elucidate Drug-Induced Metabolic Changes in a Cancer Cell Line"

### Supplementary Figures

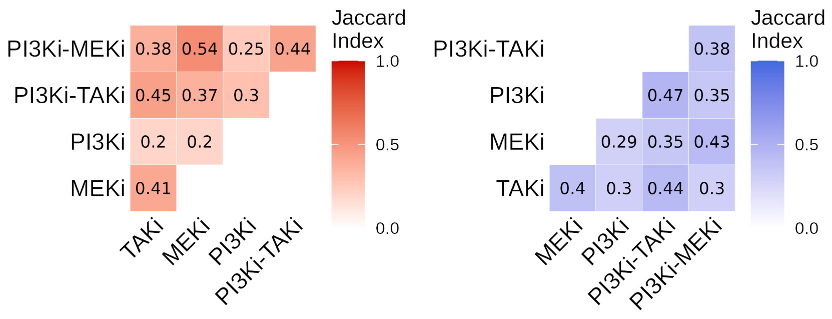


**Figure 6. Jaccard index correlation plots between up-regulated (left panel) and down-regulated (right panel) genes across treatment conditions.** Darker hues indicate higher correlation values.

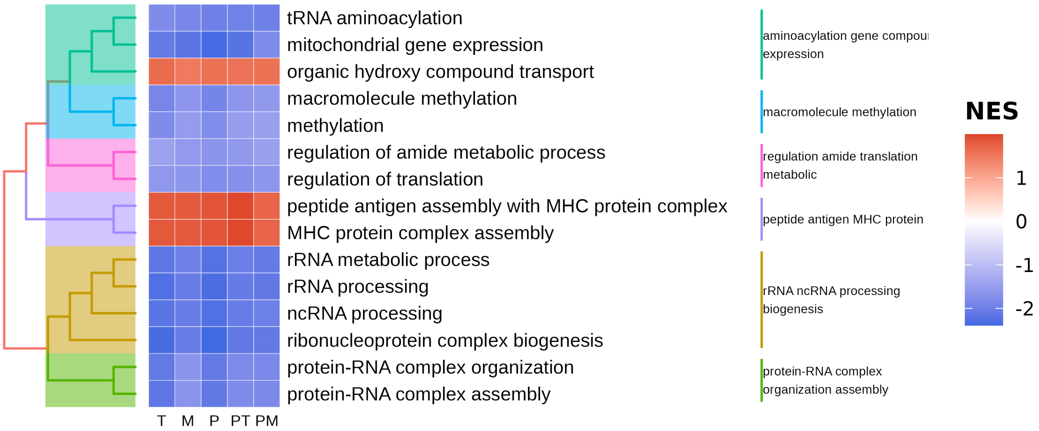


**Figure 7. Common GSEA gene sets shared across all treatment conditions.** Darker blue indicates lower, negative normalised enrichment scores (NES) values, while darker red indicates higher, positive NES values. T, M and P correspond to treatments TAKi, MEKi, and PI3Ki, respectively; PT and PM refer to combinations.

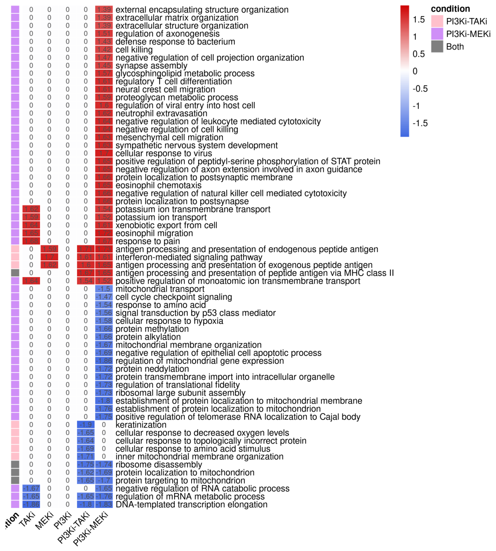


**Figure 8. GSEA gene sets significantly altered in the combinatorial treatments but not in the individual drug treatments**. Darker blue indicates lower, negative normalised enrichment scores (NES) values, while darker red indicates higher, positive NES values.

### 5. Supplementary Table Captions

**Supplementary Table 1**. Results of the differential expression analysis. Sheets labelled with the condition name show the detailed results for the corresponding condition. Columns include geneID, geneSymbol, log2FoldChange and the adjusted p-values.

**Supplementary Table 2**. Summary of the task curation process. The sheet labelled *Summary* shows the initial and final status of each task, where the status can be *optimal*, *infeasible* or *inconsistent*. Sheets *Initial_Tasks* and *Curated_Tasks* shows the full list of initial tasks from [36] and the final tasks curated for the Human 1 Model; the last sheed (Task Metadata) shows the metadata for each task, including the number of genes, the number of reactions, the number of metabolites, and the number of tasks that each gene is involved including the task ID, system, subsystem, description, source and comments.

**Supplementary Table 3**. Results of TIDE analysis. Sheet labelled *Summary* depicts the metabolic pathways affected in at least one condition, while sheets labelled with the condition name show the detailed TIDE results for the corresponding condition.

**Supplementary Table 4**. This table summarises the results of the synergy inference of the TIDE results. Sheets labelled *TIDE_synergies_PI3Ki-TAKi* and *TIDE_synergies_PI3Ki-MEKi* display the synergistic inference using the TIDE results for the combinatorial treatments PI3Ki-TAKi and PI3Ki-MEKi, respectively. Sheets labelled *TIDEessential_synergies_PI3Ki-TAKi* and *TIDEessential_synergies_PI3Ki-MEKi* display the synergistic inference using the TIDE-essential results for the combinatorial treatments PI3Ki-TAKi and PI3Ki-MEKi, respectively.

**Supplementary Table 5**. Comparison of the results obtained using TIDE and TIDE-essential. Sheet labelled *Summary* resumes the total number of altered tasks uniquely found by each algorithm and the ones in common. Sheets labelled with the condition name show the detailed TIDE results for the corresponding condition.

### Supplementary Methods

In this section, more detailed explanations of the different computational methods used in this study are provided.

*Constraint-based modeling*

Constraint-based modelling is a family of computational methods that formulate constraints as equations to define the feasible states of a metabolic system or flux space. A metabolic network can be represented using the stoichiometric matrix $N$ of $mxr$ where $m$ corresponds to the number of metabolites ($M$) and $r$ the number of reactions ($R$). The fundamental constraints used to define the flux space are the mass balance constraint:

$$N \cdot \vec{v} = 0 \qquad \text{(Eq. 3)}$$

the thermodynamic constraints on irreversible reactions

$$v_i \geq 0 \forall i \in Irrev \qquad \text{(Eq. 4)}$$

and the capacity constraints which limit the total flux through each reaction:

$$\alpha_i \leq v_i \leq \beta_i \qquad \text{(Eq. 5)}$$

When applied to boundary reactions, the capacity constraints define the metabolites produced/consumed by the system [51]. By boundary reaction, we mean reactions representing sources/sinks that consume/produce a single metabolite. Boundary reactions are referred to as exchange reactions and when defined over each metabolite present in the environment. Boundary reactions are used to define growth conditions as well as to set sources and sinks to internal metabolites that correspond to a known dead-end or gap metabolites [52].

*Flux Balanace Analyis*

Flux Balance Analysis (FBA) is a constraint-based approach for finding flux distributions that maximise the flux through the biomass reactions [53]. FBA is formulated as a Linear Program using the mass balance constraint (5), the thermodynamic constraint (5) and the boundary constraint (5). The resulting linear optimisation problem of FBA is the following:

$$\begin{aligned} &\text{Maximize:} Z = v_{biomass} \\ &\text{s.t.} \\ &\quad N \cdot \vec{v} = 0 \\ &\quad \beta_i \leq v_i \leq \beta_i \end{aligned} \qquad \text{(Eq. 6)}$$

*Parsimonious flux balance analysis*

Parsimonious Flux Balance Analysis (pFBA) is a constraint-based approach for finding flux distributions that minimise the total flux, and it is usually applied to generate solutions without inconsistent thermodynamic loops [50]. The metabolic flux for a given task $t$ is computed by minimizing the total flux subject to the constraints that defined the feasible flux space 5, 5 and 5, an approach commonly referred as parsimonious Flux Balance Analysis (see equation Eq. 7). To formulate pFBA, reversible reactions must be split into irreversible reactions in opposite directions to guarantee non-negative fluxes. After converting the network into an irreversible pFBA is formulated as follows:

$$
\begin{aligned}
&\text{Minimize:} Z = \sum_i v_i \\
&\text{s.t.} \\
&\quad N \cdot \vec{v} = 0 \\
&\quad 0 \leq v_i \leq \beta_i \\
&\quad \boldsymbol{c}^\top \boldsymbol{v} = Z^*
\end{aligned}
\qquad \text{(Eq. 7)}
$$

*Metabolic Tasks*

A metabolic task is defined as a nonzero flux through a reaction or a pathway leading to the production of metabolite P from metabolite S. In general, producing a metabolite P from a precursor S usually requires cofactors such as ATP, NADH and/or NADPH as well as the balancing of protons, water, etc.

**Table 3.** Example of a metabolic task

| Metabolites | Type | Lower bound | Upper bound |
|---|---|---|---|
| Glucose | Input | 1 | 1 |
| Oxygen | Input | 6 | 6 |
| Pi | Input | 30 | 32 |
| ADP | Input | 30 | 32 |
| $H^+$ | Input | 30 | 32 |
| ATP | Output | 30 | 32 |
| $H_20$ | Output | 36 | 38 |
| $CO_2$ | Output | 6 | 6 |

Thus, a metabolic task can more generally be defined as a set of input metabolites $M^t_{in}$ that have to be converted into a set of output metabolites $M^t_{out}$. To implement a metabolic task $t$, each input metabolite $i$ is associated with a source reaction $b^{in}_i \forall i \in M_{in}$ and each output metabolite $j$ has an associated sink $b^{out}_j \forall j \in M_{out}$. Sources and sinks are then bound according to the definition of the task (see Table 3 for an example). Finally, to calculate the flux distribution for a task $t$, we applied parsimonious Flux Balance Analysis Eq. 7 to find the feasible solution that minimises the sum of all the fluxes. This solution is usually used a the flux distribution for a task, and it is used to infer the active or flux-carrying reactions on the pathways.

*Mapping gene expression into reaction using Gene–protein–reaction rules*

Gene–protein–reaction (GPR) rules are typically represented as logical expressions combining genes using AND (for protein complexes) and OR (for isoenzymes). Each reaction usually has a GPR (see Fig. 9). Mapping values into the a GPR can be done using a recursive function that evaluatess whether a reaction is active given a gene–protein–reaction (GPR) rule and a set of active genes. The evaluation is performed recursively over the GPR expression tree, which consists of gene identifiers (leaf nodes) and logical operators such as **AND** and **OR** (internal nodes) The function EvaluateGPR traverses the GPR expression tree:

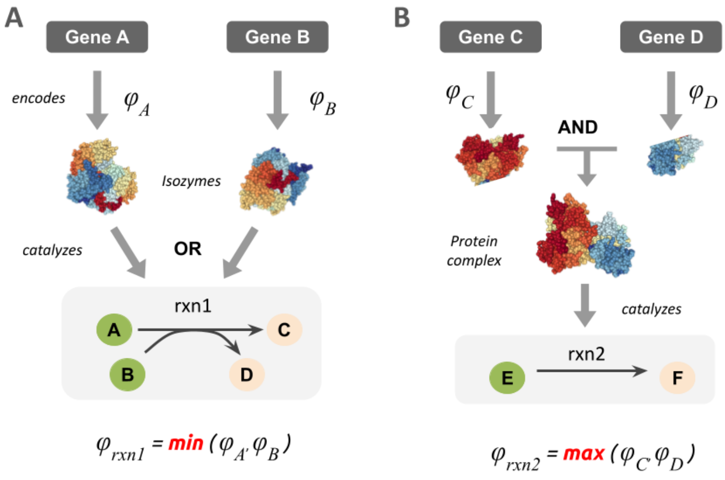


**Figure 9. Example of gene-protein-reaction-fules**. Panel A shows an example of an OR relation between genes A and B and the metabolic reaction rxn1, and corresponds to the case of two isoenzymes coding the same reaction. Panel B shows an example of an AND relation between genes C and D and the metabolic reaction rxn2, and corresponds to the case of two genes encoding subunits of an enzyme complex.

**Algorithm 1** Safe Evaluation of GPR Expression

```
function SafeEvalGPR(expr, values_dict, or_func)
    if expr is a GPR object then
        return SafeEvalGPR(expr.body, values_dict, or_func)
    else if expr is a gene name (Name) then
        if expr.id not in values_dict then
            return 0
        else
            return values_dict[expr.id]
        end if
    else if expr is a boolean operation (BoolOp) then
        if expr.op is OR then
            if or_func = "max" then
                return max_i SafeEvalGPR(expr.values[i], values_dict, or_func)
            else if or_func = "sum" then
                return ∑_i SafeEvalGPR(expr.values[i], values_dict, or_func)
            else
                raise UnsupportedGPROperator
            end if
        else if expr.op is AND then
            return min_i SafeEvalGPR(expr.values[i], values_dict, or_func)
        else
            raise TypeError("unsupported operation")
        end if
    else
        return 0
    end if
end function
```

This algorithm evaluates gene–protein–reaction (GPR) expressions based on the activity values of individual genes provided in a dictionary (`values_dict`). If the mapped gene values are continuous (e.g. gene expression or log-fold change values), the **AND** and **OR** operators are replaced by **MIN** and **MAX/SUM**, respectively. The algorithm proceeds as follows:

- If the input expression is a GPR object, its internal logical structure (i.e., its `body`) is evaluated recursively.
- If the expression is a gene (a leaf node), the algorithm returns its corresponding value from `values_dict`. If the gene is not found, a default value of `0` (inactive) is returned.
- If the expression is a Boolean operation:
  - For **OR** operators, the algorithm returns either the **maximum** or the **sum** of the child evaluations, depending on the value of the `or_func` parameter. This allows for different biological interpretations of isoenzymes—either functionally redundant (`max`) or additive (`sum`).
  - For **AND** operators, the algorithm returns the **minimum** of the child evaluations, corresponding to the requirement that all genes in an enzyme complex must be active for the reaction to be functional.
- Else the expression, the algorithm returns `0`.

For the application of the mapping for the TIDE algorithm, we used the *absolute maximum* instead of the maximum since log-Fold changes can have negative values, as described and the original paper [35].

*In-silico gene knock-outs*

*In-silico* gene knock-outs simulate the deletion of genes by constraining the associated reactions in a genome-scale metabolic model. Since genes are mapped to reactions via GPR associations, a gene deletion translates into setting the flux of all reactions catalysed exclusively by the corresponding gene product(s) to zero. In the context of FBA, this is implemented by modifying the flux bounds of the affected reactions: for each reaction $i$ associated with the knocked-out gene, we set $\alpha_i = \beta_i = 0$, effectively removing the reaction from the feasible flux space. The modified model is then solved using FBA (Eq. 6) to determine how the gene deletion affects the optimal objective value, i.e. the biomass production. A significant reduction or complete loss of biomass flux indicates that the deleted gene is essential under the given environmental conditions. This approach allows for the systematic prediction of gene essentiality and the identification of potential drug targets or synthetic lethal gene pairs *in silico*. To predict the set of essential genes of a metabolic task, we just tested for the feasibility of the problem after inactivating the reaction related to the gene being knocked; if the problem is infeasible, the tested gene is defined as essential for the task.
